# Hypoxia dampens innate immune signalling at early time points and increases Zika virus replication in iPSC-derived macrophages

**DOI:** 10.1101/2023.02.21.529434

**Authors:** Mirjam Schilling, Alun Vaughan-Jackson, William James, Jane A McKeating

## Abstract

Type I interferons (IFNs) are the major host defence against viral infection and are induced following activation of cell surface or intracellular pattern recognition receptors, including retinoic-acid-inducible gene I (RIGI)-like receptors (RLRs). All cellular processes are shaped by the microenvironment and one important factor is the local oxygen tension. The majority of published studies on IFN signalling are conducted under atmospheric (18%) oxygen conditions, that do not reflect the physiological oxygen levels in most organs (1-5% O_2_). We studied the effect of low oxygen on IFN induction and signalling in induced Pluripotent Stem Cell (iPSC)-derived macrophages as a model for tissue-resident macrophages and assessed the consequence for Zika virus (ZIKV) replication. Hypoxic conditions dampened the expression of interferon-stimulated genes (ISGs) following RLR stimulation or IFN treatment at early time points. RNA-sequencing and bio-informatic analysis uncovered several pathways including changes in transcription factor availability, the presence of HIF binding sites in promoter regions, and CpG content that may contribute to the reduced ISG expression. Importantly, hypoxic conditions increased ZIKV replication at early time points, emphasizing the importance of understanding how low oxygen conditions in the local microenvironment affect pathogen sensing and host defence.

## Introduction

Successful virus infection is dependent on the presence of specific cellular host factors and simultaneously needs to evade both innate and adaptive immune responses. The cellular microenvironment can affect a multitude of pathways that impact virus replication. Recent studies from our laboratory have identified a role for oxygen tension in regulating cellular susceptibility to virus infection (reviewed in (Liu et al. 2020)). Depending on the blood supply and metabolic demand the oxygen tension can vary between 1% and 5% in different organs (Carreau et al. 2011). This contrasts with many reported studies that employ *in vitro* tissue culture model systems that are maintained at 18% atmospheric oxygen. Lower oxygen conditions, referred to as hypoxia, will inactivate multiple oxygen sensing mechanisms, including the prolyl hydroxylases (PHD) and factor inhibiting HIF (FIH)-hypoxia inducible factor (HIF) pathways that stabilise HIF expression (Taylor and Scholz 2022, Pugh and Ratcliffe 2017). Hypoxia can modulate the replication of a wide number of viruses (Liu et al. 2020), where HIFs enhance the replication of hepatitis B (Wing, Liu, et al. 2021) and Epstein Barr viruses (Jiang et al. 2006, Kraus et al. 2017) via direct binding to their viral DNA genomes. In contrast, a hypoxic environment suppresses influenza A virus (Zhao et al. 2020), SARS-CoV-2 (Wing, Keeley, et al. 2021, Wing et al. 2022) and HIV-1 (Zhuang et al. 2020) replication. These differing outcomes may reflect variable oxygen levels at the site of virus replication.

The type I IFN system is a crucial first line of defence to combat pathogens (McNab et al. 2015). Pathogen associated molecular patterns (PAMPs) are recognized by cellular sensors, such as Toll like receptors (TLRs), RIG-I like receptors (RLRs) or cytoplasmic DNA receptors. Through downstream activation of kinases and transcription factors, IFNs are induced and stimulate the expression of interferon-stimulated genes (ISGs) that create an antiviral state. A recent study reported that hypoxia downregulates the RLR dependent type I IFN pathway in cancer cell lines and this associated with a decreased accessibility of STAT1 and IRF3 motifs in the host chromatin (Miar et al. 2020). Since RLR signalling is a key inducer of innate immune responses we chose to study the consequences of low oxygen on IFN induction and signalling in primary immune cells. We found that hypoxia dampened ISG expression in human induced pluripotent stem cell (iPSC) derived macrophages early after stimulation with RLR activators and IFN. RNAseq analysis suggests this is mediated by a dysregulation of pathways that affect the expression and stability of ISG transcripts, including changes in transcription factor availability, the presence of HIF binding sites in promoter regions, and CpG content.

RIG-I is the main sensor that detects Zika virus (ZIKV) (Hamel et al. 2015, Lazear et al. 2016, Chazal et al. 2018, Hertzog et al. 2018, Esser-Nobis et al. 2019, Schilling et al. 2020), a member of the Flaviviridae family, that was first isolated from sentinel rhesus macaque in the Zika Forest in Uganda and later from Aedes africanus mosquitoes (Dick, Kitchen, and Haddow 1952). Despite causing a self-limiting acute febrile illness in adults, ZIKV infection during the first trimester of pregnancy is associated with multiple neurodevelopmental defects, including microcephaly in newborns (Faria et al. 2016). ZIKV has been reported to infect Hofbauer cells, macrophages within the placenta (Simoni et al. 2017) and we selected iPSC-derived macrophages as a model to study the effect of hypoxia on ZIKV infection (Buchrieser, James, and Moore 2017).

## Material and Methods

### Cells and reagents

The iPSC-derived macrophages were differentiated from the human iPSC line OX1-61 (Buchrieser et al. 2018, Vaughan-Jackson et al. 2021) and cultured in advanced DMEM/F12 supplemented with 1% penicillin/streptomycin, Glutamax (2 mM), stabilized Insulin (5 μg/ml), HEPES pH 7.4 (15 mM), M-CSF (100 ng/ml). The human iPSC derived lines used in this study are SFC841-03-01 (Dafinca et al. 2016) (donor #1) and SFC840-03-03 (Fernandes et al. 2016) (donor #2).

| *Reagent* | *Source* | *Catalogue number* |
| --- | --- | --- |
| Poly(I:C) | Sigma-Aldrich | P1530 |
| 5'pppRNA | InvivoGen | tlrl-3prna |
| IFNα2a | PeproTech | 300-02AA |

### Viruses

The Brazilian ZIKV isolate ZIKV/H. sapiens/Brazil/PE243/2015 was originally described in (Donald et al. 2016) and was propagated in Vero cells. Infectivity of viral stocks was determined by plaque assay using A549 BVDV NPro cells. These cells stably express the NPro protein of bovine viral diarrhea virus (BVDV), which induces degradation of IRF3, and are optimized for virus growth (Hilton et al. 2006).

### Cell viability assay

Viability was assessed using the CellTiter-Fluor Cell Viability Assay (Promega, G6080) according to manufacturer’s instructions. Fluorescence was measured on the Spectramax M5 plate reader using SoftmaxPro software version 5.

### qRT-PCR

Cells were lysed and total RNA extracted using the RNeasy kit (Qiagen) according to the manufacturer’s instructions. Equal amounts of cDNA were synthesized using the High Capacity cDNA Kit (Applied Biosystems) and mRNA expression determined using Fast SYBR master mix in a StepOne thermocycler (Applied Biosystems). C_T_ values were normalized to TBP (ΔC_T_). SYBR green primer probes used include TBP (for: CCCATGACTCCCATGACC, rev: TTTACAACCAAGATTCACTGTGG), NDRG1 (for: TTTGATGTCCAGGAGCAGGA, rev: ATGCCGATGTCATGGTAGGT), MX1 (for: GGCTGTTTACCAGACTCCGACA, rev: CACAAAGCCTGGCAGCTCTCTA), DDX58 (for: CACCTCAGTTGCTGATGAAGGC, rev: GTCAGAAGGAAGCACTTGCTACC) and ZIKV (for: TCGTTGCCCAACACAAG, rev: CCACTAATGTTCTTTTGCAGACAT).

### Western Blotting

Cells were lysed in RIPA buffer (20 mM Tris pH 7.5, 2 mM EDTA, 150 mM NaCl, 1% NP40, 1% SDS, 0.1% TritonX100, 0.25% Na-Deoxychalate, plus protease inhibitor) and the samples were incubated at 95 °C for 5 min. Protein lysates were separated on 10% SDS-PAGE gels and transferred onto polyvinylidene difluoride (PVDF) membranes. As primary antibodies we used anti-Mx (mouse, M143, (Flohr et al. 1999)), anti-RIG-I (mouse, AdipoGen), and anti-beta-actin (mouse, AC-15, Sigma-Aldrich). Primary antibodies were detected with peroxidase-conjugated secondary antibodies (GE Healthcare).

### Flow Cytometry Analysis

To measure cell viability, cells were stained with Live Dead Aqua (Life Technologies, UK) in addition to the primary antibody (IFNAR1, EP899Y, rabbit, abcam). After washing, cells were stained with secondary antibody (goat anti-rabbit, Alexa Fluor 633) and resuspend in BD Cell FIX. Samples were acquired on a Cyan ADP flow cytometer (Beckman Coulter) and data analysed using FlowJo (TreeStar).

### RNA-Seq and bio-informatic analysis

RNA was isolated using the RNeasy kit (Qiagen) and RNA-sequencing performed by Novogene: RNA purity was assessed with a NanoDrop 2000 spectrophotometer (Thermo Fisher Scientific) and integrity determined using a 2100 Bioanalyzer Instrument (Agilent Technologies). Sequence adapters, reads containing ploy-N and low quality reads were removed from the raw data through in-house perl scripts. The reads were mapped against the reference human genome (GRCh38/hg38) using hisat2 v2.0.5. Counts per gene were calculated using featureCounts v1.5.0-p3 and the FPKM of each gene was calculated based on the length of the gene and reads count mapped to this gene. Reads were analyzed by edgeR v3.22.05. The P values were adjusted using the Benjamini & Hochberg method. Corrected P-value of 0.05 and absolute foldchange of 2 were set as the threshold for significant changes in gene expression.

Gene set enrichment analysis was performed using GSEA_4.1.0, and hallmark gene sets retrieved from the Molecular Signatures Database (MSigDB) (Subramanian et al. 2005, Liberzon et al. 2015). As permutation-type we used “gene_set”, and as cut-off an FDR of 0.25. GraphPad Prism (version 9.3.1, GraphPad, San Diego, CA, USA) was used to plot the normalized enrichment score (NES) and colour-code by the false discovery rate (FDR). Transcription factor binding sites of HIF1A (ranging from −1000 to 100bp relative to TSS, with cut-off p=0.001) were analysed using https://epd.epfl.ch//index.php. Transcription factor enrichment analysis (TFEA) was performed through the ChEA3 background database (https://maayanlab.cloud/chea3/) to compare discrete query gene sets with libraries of target gene sets assembled from multiple orthogonal ‘omics’ datasets. (Keenan et al. 2019).

Dinucleotides were counted using R (version 4.0.3 (2020-10-10)) in Rstudio (Version 1.4.1103) to download sequences from the NCBI database (Genbank), count dinucleotides for all 16 combinations and normalizing the counts to the length of the respective sequence (R Core Team). CpG content of either the up- and downregulated ISGs, or the top 30 up- and downregulated protein-coding genes based on log2-fold or adjusted p-value were plotted with GraphPad Prism (version 9.3.1, GraphPad, San Diego, CA, USA) and a nonparametric t-test (Mann-Whitney) was performed, *p<0.05, **p<0.001.

### Quantification and statistical analysis

Data were analysed using GraphPad Prism version 9.3.1 (GraphPad, San Diego, CA, USA). All data are presented as mean values□±□SEM. Significance values are indicated as *p< 0.05; **p< 0.01; ***p< 0.001; ****p< 0.0001. n.s. denotes non-significant. Please see individual figure legends for further details.

## Results

### Hypoxia dampens ISG, but not IFNAR1 cell surface expression

iPSC-derived macrophages were differentiated from pluripotent stem cells via embryoid body intermediates as previously reported (Vaughan-Jackson et al. 2021). This protocol uses open-source media components that result in a lower basal expression of ISGs compared to cells differentiated using earlier methods, while being more responsive to inflammatory stimulation. We used independent differentiations of stem cells from the same donor, or a second, independent donor.

To investigate the effect of hypoxia on RIG-I like receptor (RLR) signalling iPSC-derived macrophages were cultured at 1% O_2_ or 18% O_2_ for 24h and transfected with Poly(I:C) (Figure 1A). Poly(I:C) is a synthetic mimetic of a double-stranded RNA that is detected by TLRs or, when transfected, by RLRs. We selected a concentration of Poly(I:C) that did not affect cell viability and induced ISG expression (Figure 1, Suppl. Figure). We measured expression of MX Dynamin Like GTPase 1 (*MX1*), an ISG solely induced by type I and type III IFNs, and *RIG-I*, which can be directly stimulated by Interferon regulatory factor 3 (IRF3) activation. As a control we measured transcript levels of the HIF regulated gene N-Myc Downstream Regulated 1 (*NDRG1*) (Figure 1A/C/E). While hypoxia induced *NDRG1* gene expression, we observed a reduction in *MX1* and *RIG-I* transcripts. Due to limiting cell availability we studied cells at 4h post Poly(I:C) treatment from an independent differentiation along with cells from an additional donor and showed a two-fold reduction in *MX1* gene expression (Figure 1B). To specifically stimulate RIG-I the cells were transfected with 5’pppRNA (Schlee and Hartmann 2010) and we observed a reduction in *MX1* and *RIG-I* gene expression at the peak of gene induction (Figure 1C). Repeat experiments showed a significant 2-4 fold reduction in *MX1* and *RIG-I* transcripts (Figure 1D). To differentiate between the effects on intracellular RNA sensing and secondary intercellular IFN signalling, we treated the iPSC-derived macrophages with IFNα2a and observed a reduction in *MX1* and *RIG-I* gene expression at 4h post-stimulation (Figure 1E/F). This reduction in ISG expression waned at later time points, similar to our observations after Poly(I:C) transfection. The hypoxic suppression of *MX1* and *RIG-I* transcripts following IFN treatment (24h) was more pronounced at the protein level, with Mx protein expression reduced 7-to 11-fold (Figure 1G).

**Fig. 1:**
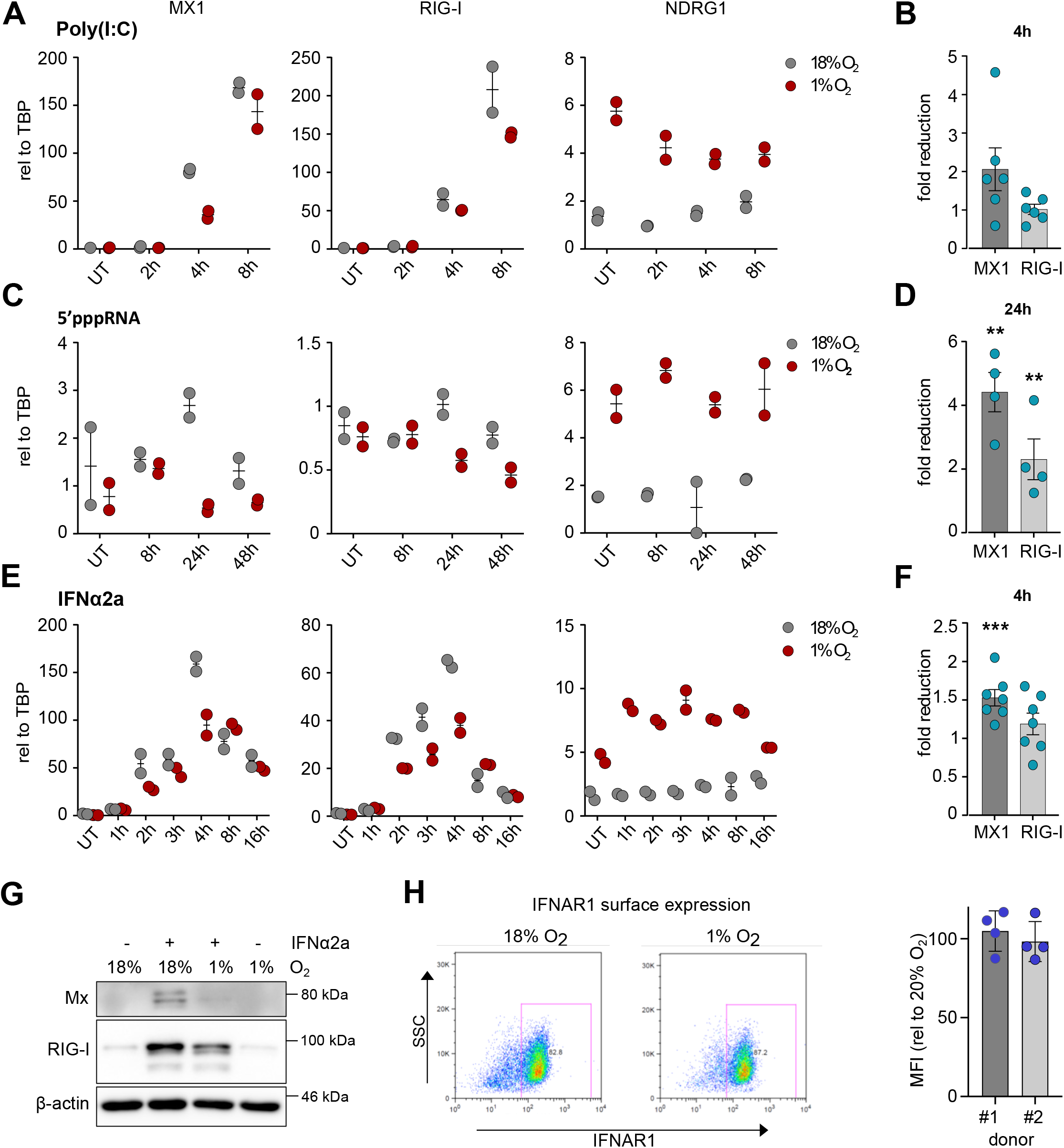
Hypoxia dampens ISGs but not IFNAR1 cell surface expression. RT-qPCR analysis of iPS-derived macrophages **A)** transfected with Poly(I:C) (0.2 μg/ml) 24h after incubating under normoxic (18% oxygen) or hypoxic (1% oxygen) conditions. One representative experiment of donor #1 with two technical replicates is shown. **B)** Two independent differentiations of donor #1 with n=4 independent experiments, plus n=2 experiments with donor #2 relative to normoxia are shown for one timepoint. **C)** transfected with 5’pppRNA (0.2μg/ml) 24h after incubating under normoxia or hypoxia. One representative experiment of donor #1 with two technical replicates is shown. **D)** One independent differentiation of donor #1 with n=2 and donor #2 with n=2 relative to normoxia is shown for one timepoint. **E)** treated with IFNα2a (100U/ml) 24h after exposure to normoxia or hypoxia. One representative experiment of donor #1 with two technical replicates is shown. F) Two independent differentiations of donor #1 with n=5, plus n=2 experiments with donor #2. Unpaired t-test with *p<0.05, **p<0.01., ***p<0.001. **G)** Western Blot analysis of donor#1 exposed to normoxia or hypoxia for 24h followed by a 24h treatment with IFNα2a (100U/ml). One representative blot (n=3) is shown. **H)** Flow cytometric analysis of IFNAR1 cell surface expression after 24h of hypoxia. One representative FACS plot is shown for donor#1 and the MFI normalized to 18% for n=3 independent experiments for both donors.

One explanation for the observed dampening in ISG expression following Poly(I:C) and 5’pppRNA treatment may be a reduction in Interferon Alpha And Beta Receptor Subunit 1 (IFNAR1) expression. To examine this, we stained IFNAR1 on iPSC-derived macrophages cultured at 18% or 1% oxygen for 24h. IFNAR1 surface expression was unchanged in hypoxic conditions compared to normoxia (Figure 1H and Suppl Figure 2). Overall, these findings show that changes in the local oxygen tension affect RIG-I like receptor sensing and IFN signalling in iPSC-derived macrophages that result in a dampened ISG response.

### Transcriptomic analysis of IFN-treated hypoxic iPSC-derived macrophages reveals a broad dampening of IFNα and IFNγ hallmark genes

To investigate whether hypoxia leads to global reduction in ISGs other than Mx1 and RIG-I we performed a RNAseq analysis of IFN-treated iPSC-derived macrophages 4h post-stimulation with IFNa2a. A total of 2174 genes were up-, and 1854 genes downregulated in hypoxia compared to normoxia (Figure 2A). Gene set enrichment analysis (GSEA) showed a significant increase in genes connected to hypoxia and glycolysis confirming the cellular response to low oxygen (Figure 2B/C). Additional pathways related to MTORC, TNFα, IL-2, Notch or Kras signalling were significantly increased. More importantly, GSEA confirmed that hypoxia reduced IFNα and IFNγ hallmark genes (Figure 2C). We noted that not all of the 103 differentially expressed ISGs were suppressed under the hypoxic conditions (Figure 2D). We found a total of 29 upregulated and 74 downregulated ISGs. Interestingly, many of the well-described direct anti-viral acting ISGs, such as *MX1*, Interferon-Induced Protein With Tetratricopeptide Repeats 1 (*IFIT1*) or SAM And HD Domain Containing Deoxynucleoside Triphosphate Triphosphohydrolase 1 (*SAMHD1*) were suppressed under the hypoxic conditions.

**Fig. 2:**
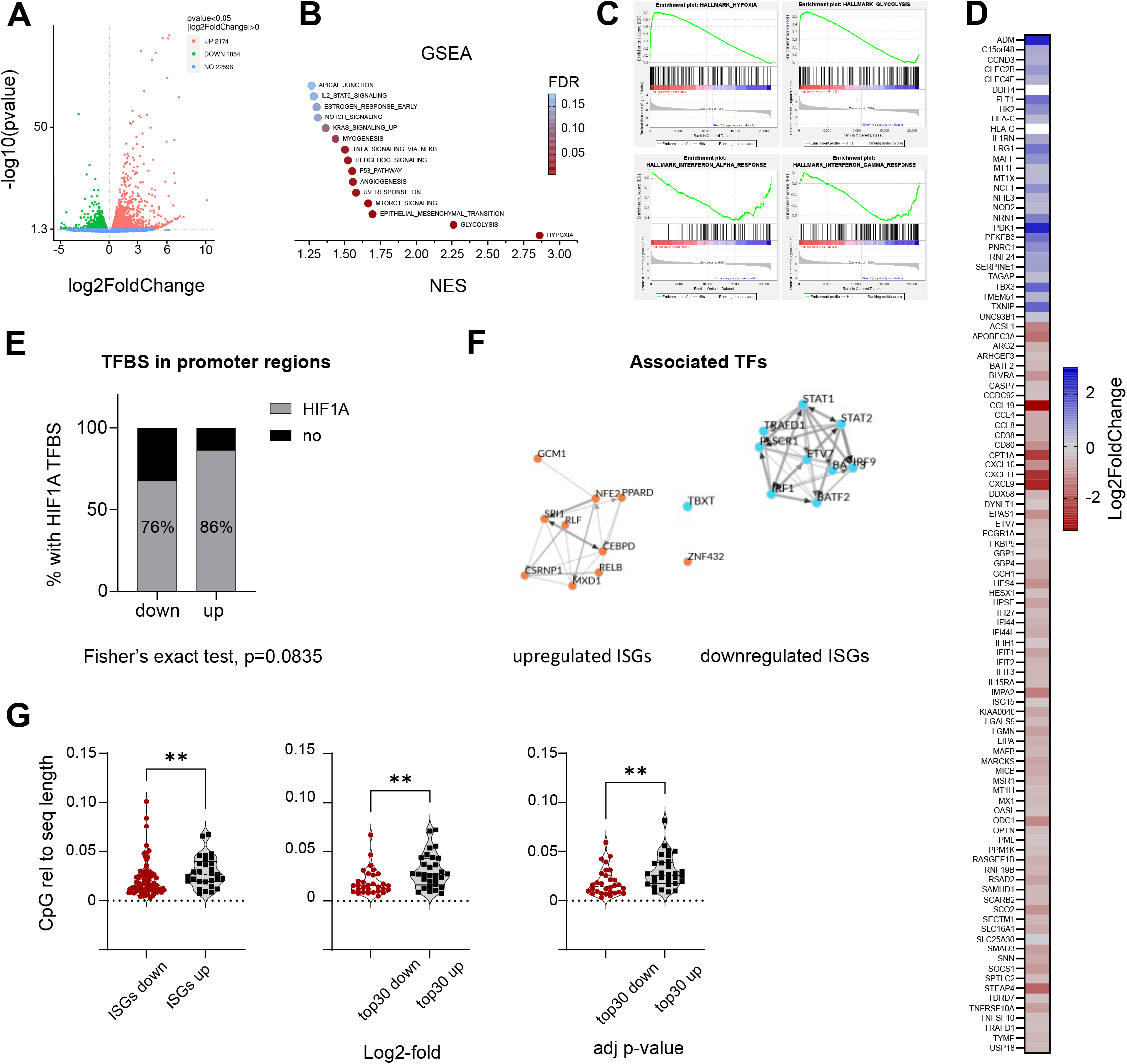
RNAseq analysis of IFN-treated iPS-derived macrophages in hypoxia reveals a broad dampening of IFNα and IFNγ hallmark genes. **A)** Volcano plot showing the distribution of differentially expressed genes between cells in hypoxia vs normoxia of donor #1 4h after IFNα2a stimulation. **B)** Gene set enrichment analysis using the hallmark gene sets from the molecular signature database (MSigDB). Plots show the normalized enrichment score (NES) and colour-coded by the false discovery rate (FDR). **C)** GSEA shows positive enrichment scores of gene sets associated with hypoxia or glycolysis, and negative enrichment scores of gene sets associated with IFNα or IFNγ responses. **D)** Heatmap of 103 differentially expressed ISGs. **E)** Bar graph representing the percentage of up- or downregulated ISGs with a ARNT::HIF1A transcription factor binding motif in their promoter region (−1000 relative to transcription start site). **F)** Transcription factor (TF) enrichment analysis of TFs associated with up- or downregulated ISGs. Shown is the local network of the top 10 TFs ranked across libraries. **G)** CpG content relative to sequence length of the transcript of either the up- and downregulated ISGs, or the top 30 up- and downregulated protein-coding genes based on log2-fold or adjusted p-value. Nonparametric t-test (Mann-Whitney), *p<0.05, **p<0.01.

To investigate the mechanisms through which hypoxia affects ISG expression, we sought to identify pathways that could discriminate between the differentially expressed ISGs. We first analysed the presence or absence of transcription factor binding motifs in the respective promoter regions (−1000 relative to transcription start site) using the eukaryotic promoter database. Interestingly, the majority of promoter regions analysed encoded hypoxic response elements (HREs) (Figure 2E). However, only about 10% more of the upregulated ISGs encoded a HIF binding motif compared to the downregulated ISGs, suggesting this is unlikely to account for the differences in transcriptional regulation between the groups.

To gain a deeper insight into the transcriptional regulation of the ISGs we performed a transcription factor (TF) enrichment analysis to identify factors that associate with up- or down-regulated ISGs using the ChIP-X Enrichment Analysis 3 (ChEA3) (Keenan et al. 2019). The predicted TFs associated with up- and downregulated ISGs are shown in Figure 2F. We asked whether any of these key transcriptional regulators were differentially expressed under hypoxic conditions to account for the variable ISG expression. Comparing the predicted TFs encoded in the up- or downregulated genes identified several TFs associated with the upregulated ISGs, including CSRNP1, MXD1, PPARD, RLF and ZNF432. Additionally, the expression of BATF2, a TF associated with the downregulated group of ISGs was decreased. These data suggest that hypoxia causes an imbalance in the expression of several TFs that will impact the regulation of ISG subsets.

The level of transcripts in a cell is, however, not only affected by the expression or binding of transcription factors, but also by the availability of nucleotides. Our RNAseq analysis showed that hypoxia affects nucleoside metabolism (Suppl Figure 3). This is in line with a reported depletion of nucleotides under hypoxia and an increase in the assembly of the multienzyme purinosome complex (Cohen, Geck, and Toker 2020, Doigneaux et al. 2020). Interestingly, Shaw et al. reported that ISGs have a lower CpG content than the human transcriptome (Shaw et al. 2021). We explored whether an imbalance in the pool of nucleosides could account for the increase and decrease of certain genes under hypoxia. To investigate this hypothesis, we quantified the dinucleotide content of our two ISG groups and found they differed significantly in their CpG content (Figure 2G). To investigate whether this is a common pathway of ISG and host gene regulation under hypoxia we analysed the CpG content of the 30 most highly up- or downregulated genes in our dataset, based on either their 2log fold or adjusted p-values (Figure 2G). Interestingly, not only the up- and downregulated ISGs but also the 30 top hypoxic up- or downregulated host genes differed in their CpG content. This suggests that hypoxia may affect gene expression based on the dinucleotide content of transcripts.

Overall, our data show that hypoxia most likely affects the expression of ISGs through changes in transcription factor availability, the presence of HIF binding sites in promoter regions, and intrinsic properties of the transcripts, such as CpG content.

### Hypoxia promotes ZIKV replication in iPSC-derived macrophages

Since our RNAseq analysis showed a robust suppression of IFNα and IFNγ hallmark genes, we speculated this would associate with an increase in virus replication. We chose to study ZIKV as it is mainly sensed by RIG-I (Hamel et al. 2015, Lazear et al. 2016, Chazal et al. 2018, Hertzog et al. 2018, Esser-Nobis et al. 2019, Schilling et al. 2020). To investigate the functional consequences of a dampened ISG response under hypoxia, we infected iPSC-derived macrophages kept in hypoxia or normoxia with ZIKV (MOI 1) and measured viral RNA levels 24h after infection by RT-qPCR. We detected increased ZIKV RNA in the hypoxic iPSC-derived macrophages (Figure 3A). We next wanted to test whether ZIKV replication is promoted through an increase in cell entry or at a post-entry step. We therefore omitted priming cells with hypoxia for 24h before infection and only applied low oxygen conditions after infection. Interestingly, we still observed an increase in ZIKV replication in the non-primed cells under hypoxia (Figure 3B), consistent with a model where low oxygen conditions promote ZIKV replication at post-entry steps.

**Fig. 3:**
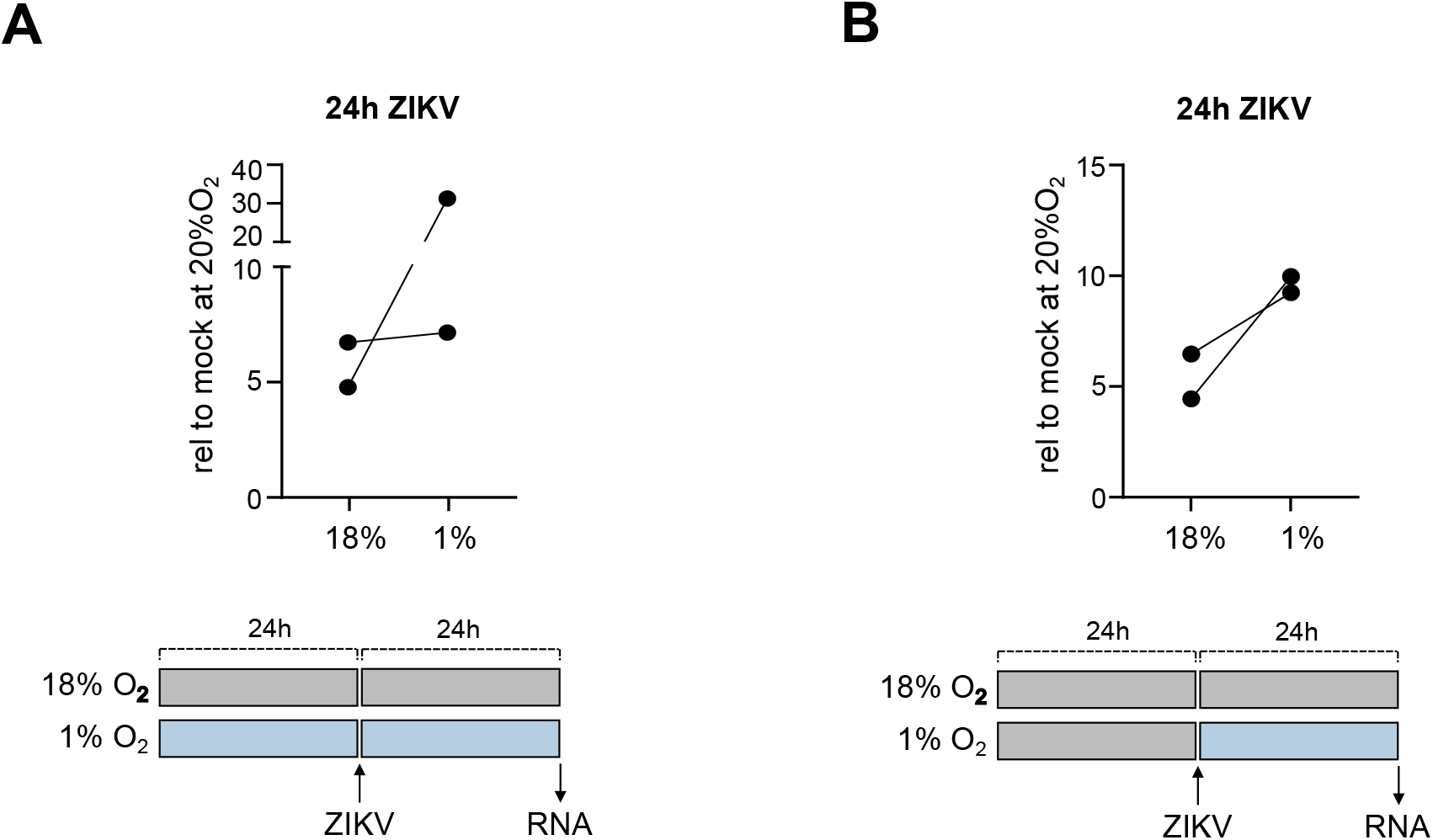
Hypoxia promotes ZIKV replication in iPS-derived macrophages. **A)** Comparison of ZIKV RNA levels in iPS-derived macrophages primed with 18% or 1% O_2_ for 24h before infection with ZIKV (MOI 1). **B)** Comparison of iPS-derived macrophages cultured at 18% O_2_ before infection and propagating at either 18% or 1% O_2_ after infection with ZIKV (MOI 1). RT-qPCR for ZIKV RNA 24h after infection, n=2 of one differentiation of donor #1.

## Discussion

Our study shows that hypoxia dampens ISG expression in iPSC-derived macrophages at early times following stimulation, that are important in the initiation of rapid antiviral responses. In line with this, we detected an increase in ZIKV replication under hypoxia. Interestingly, we observed the most significant dampening of ISG expression after stimulation with IFNα2a, suggesting that both PAMP sensing as well as interferon signalling are affected by the changes in the oxygen tension. It is tempting to speculate that the higher fold-difference we detected in the sensing of 5’ppp RNA compared to IFN signalling reflects an additive effect of hypoxia on both IFN induction and IFN signalling. Our findings are consistent with a recent report describing that hypoxia suppresses the induction of type I IFN in monocytes (Peng et al. 2021). Furthermore, our results agree with earlier studies, reporting an increase in the replication of other members of the *Flaviviridae* family, such as hepatitis C and dengue viruses under low oxygen conditions (Frakolaki et al. 2018, Vassilaki et al. 2013, Wilson et al. 2012).

Our findings may be relevant in the context of ZIKV infection *in vivo*, where oxygen levels in the placenta are approx 2-3% in the first trimester and 5-8% in the second and third trimesters of pregnancy (Tuuli, Longtine, and Nelson 2011). Our data suggest these conditions could contribute to the severe ZIKV-induced neurodevelopmental defects due to lower innate immune responses and higher ZIKV replication. Additionally, in-vivo experiments in mice show an increased permeability of the blood brain barrier 6h after hypoxia, which could increase virus replication in the brain (Engelhardt, Patkar, and Ogunshola 2014).

Hypoxia was shown to reduce host responses to TLR ligands in airway epithelia cells (Polke et al. 2017) in a HIF1α-dependent manner. However, our data suggest that the presence or absence of HIF-binding sites in the promoter regions of ISGs does not associate with the overall up- or downregulation observed. We recognise that bio-informatic predictions have significant limitations and this is reinforced by a recent HIF ChIP-seq analysis that identified only 500-100 binding sites in the human genome (500-1000), despite their high abundance (over 1 million) (Kindrick and Mole 2020, Schödel et al. 2011). Apart from HIF-dependent changes other cellular pathways are activated under hypoxic conditions and will affect the transcriptome. Miar and colleagues reported a minimal role for HIF-1α or HIF-2α in regulating the hypoxic suppression of ISGs in cancer cell lines, but decreased chromatin accessibility of genomic regions relevant for type I IFN pathways (Miar et al. 2020). Recent studies reported that lactate, a glycolytic product that is increased during hypoxia, regulates gene transcription via lactylation of histones (Xin et al. 2022, Zhang, Tang, et al. 2019). In tumor-infiltrating myeloid cells H3K18 lactylation increased N6-adenosine-methyltransferase 70 kDa subunit (*Mettl3*) expression that modified *Jak1* mRNA, thereby strengthening the immunosuppressive functions (Xiong et al. 2022). Interestingly, lactate inhibits oligomerization of MAVS, the adapter molecule for RLRs, and dampens RLR signalling (Zhang, Wang, et al. 2019). The detailed mechanisms through which hypoxia affects IFN signalling in the iPSC macrophages will require further study. Our data suggest that hypoxia affects ISG expression by the disturbance of multiple pathways.

The increasing relevance to study arthropod-borne viruses due to the changing geographical location and transmission patterns associated with global warming, emphasises the importance to discover new therapeutic targets that may be broadly effective against diverse members of the *Flaviviridae* family (Ryan et al. 2019). In addition, virus infections and associated inflammation can affect local oxygen levels. Mitochondrial damage will increase reactive oxygen species, leading to a change in local metabolite availability and oxygen consumption (Daniels et al. 2019, Williams, Sitole, and Meyer 2017, Li et al. 2017). Therapeutically targeting the pathways that dampen ISG expression could help treat a broad range of viral infections.

## Limitations of the study

The sample size of this study was limited by the production capacity of the embryoid body intermediates, as well as lab closures during the SARS-CoV-2 pandemic.

## Author contributions

Conceptualization: M.S., J.MK. Methodology: M.S., A.VJ., J.MK., W.J. Investigation: M.S. Formal analysis and visualisation: M.S. Resources: M.S., A.VJ., J.MK., W.J. Writing – original draft preparation: M.S. Writing – review and editing: all authors.

## Acknowledgements

We would like to thank Jan Rehwinkel for the kind gift of the anti-Mx (mouse, M143, (Flohr et al. 1999)) and anti-RIG-I (mouse, AdipoGen) antibody.

---

**Suppl. Fig. 1: Poly(I:C) treatment does not affect cell viability.**

iPS-derived macrophages were treated with 0.2ug/ml Poly(I:C) for 3 days and live-cell protease activities measured using the GF-AFC Substrate (live-cell protease substrate), MultiTox-Fluor Multiplex Cytotoxicity Assay (G9201, Promega). Data represent the AFC fluorescence (RFU) expressed relative to untreated cells.

**Suppl. Fig. 2: Gating strategy to stain for IFNAR1 surface expression on iPS-derived macrophages.**

After 24h in 1% or 18% O_2_ iPS-derived macrophages were stained with Live Dead Aqua and IFNAR1 antibody. FACS plots shown are a representative example of the gating strategy used in each experiment

**Suppl. Fig. 3: Hypoxia dysregulates nucleoside metabolism.**

Gene Ontology enrichment analysis scatter plot depicting the most significant 30 terms enriched under hypoxia for donor#1. Figure provided by novogene.

