## Supplementary material for "Hypoxia dampens innate immune signalling at early time points and increases Zika virus replication in iPSC-derived macrophages": Suppl Figures

### Supplementary Figures

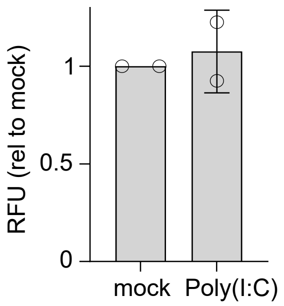


**Suppl. Fig. 1: Poly(I:C) treatment does not affect cell viability.**

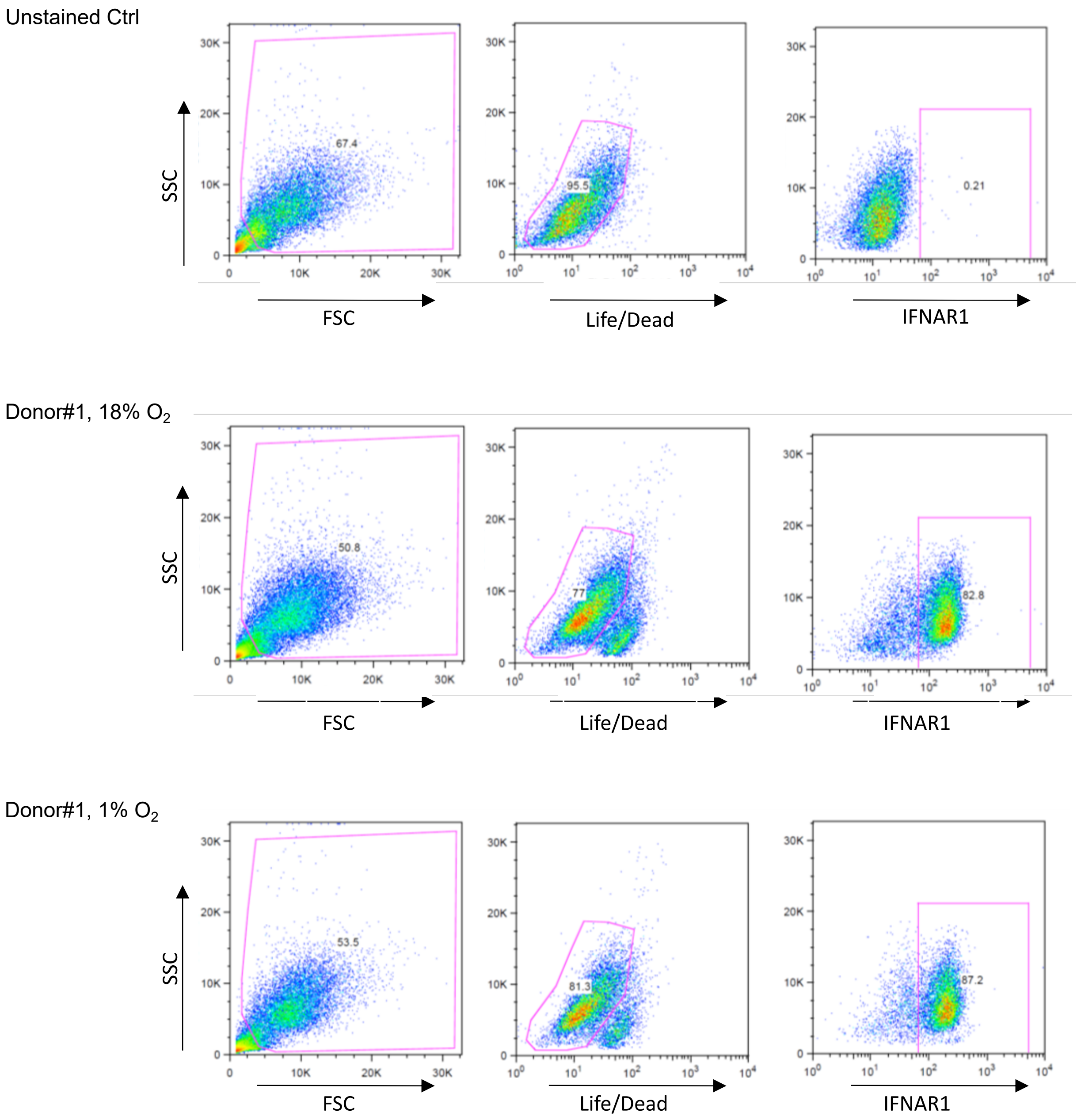


**Suppl. Fig. 2: Gating strategy to stain for IFNAR1 surface expression on iPSC-derived macrophages.**

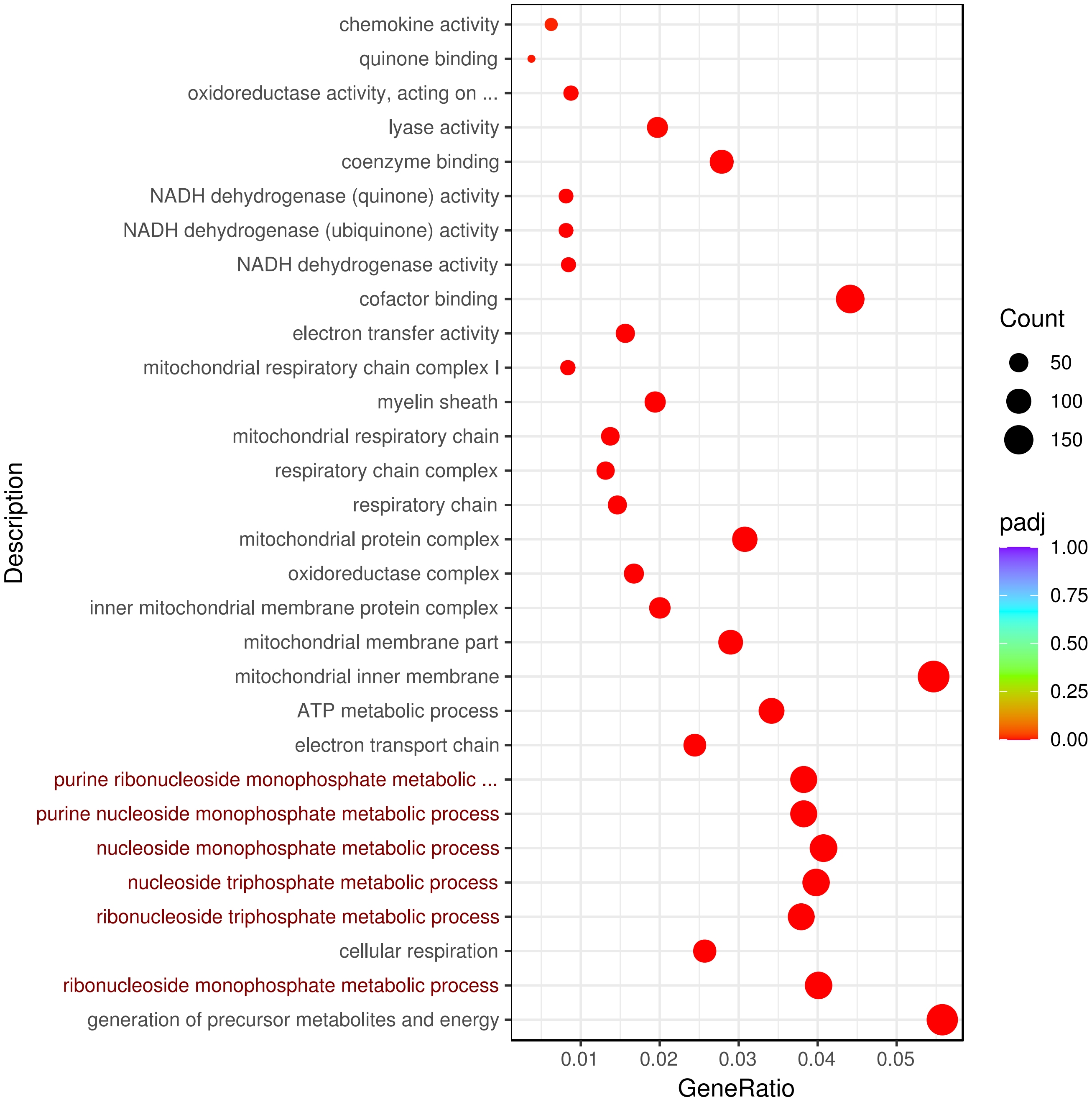


**Suppl. Fig. 3: Hypoxia dysregulates nucleoside metabolism.** .
